# The chromatin-regulating CoREST complex is animal specific and essential for development in the cnidarian *Nematostella vectensis*

**DOI:** 10.1101/2021.11.11.468220

**Authors:** James M Gahan, Maria Hernandez-Valladares, Fabian Rentzsch

**Affiliations:** Sars International Centre for Marine Molecular Biology, University of Bergen, Thormøhlensgate 55, 5006 Bergen, Norway; Proteomics Facility of the University of Bergen (PROBE), University of Bergen, 5020 Bergen, Norway; Department for Biological Sciences, University of Bergen, Thormøhlensgate 53, 5006 Bergen, Norway

## Abstract

Chromatin-modifying proteins are key players in the regulation of development and cell differentiation in animals. Many individual chromatin modifiers, however, predate the evolution of animal multicellularity and how they became integrated into the regulatory networks underlying development is unclear. Here we show that CoREST is an animal-specific protein that assembles a conserved, vertebrate-like histone-modifying complex including Lsd1 and HDAC1/2 in the sea anemone *Nematostella vectensis.* We further show that NvCoREST expression overlaps fully with that of NvLsd1 throughout development. *NvCoREST* mutants, generated using CRISPR-Cas9, reveal essential roles during development and for the differentiation of cnidocytes, thereby phenocopying *NvLsd1* mutants. We also show that this requirement is cell autonomous using a cell-type-specific rescue approach. Together, this shows that the evolution of CoREST allowed the formation of a chromatin-modifying complex that was present before the last common cnidarian-bilaterian ancestor and thus represents an ancient component of the animal developmental toolkit.

## Introduction

Understanding the evolution of animal development and cell differentiation requires analysis of the gene regulatory programs that direct these processes. In recent years, comparisons between animal groups have found a remarkable degree of conservation of transcription factors and signaling pathways throughout the animal kingdom (*1–6*). This suggests that changes in the repertoire of these genes alone cannot explain the diversification of development processes. Regulation of chromatin has been shown to be another essential aspect of transcription during development (*7–9*), but its potential role in the evolution of developmental gene regulation has received little attention.

Many chromatin regulators are ancient and predate the evolution of animal multicellularity. One possible way to evolve roles in developmental programs is the integration of chromatin regulators into multiprotein complexes that facilitate the coordination of different regulatory (e.g., enzymatic) activities and/or facilitate targeting to specific genomic loci. Here, we use the CoREST complex to explore this scenario in an early-diverging group of animals.

The CoREST complex was initially discovered in mammals as a complex required for repression of neuronal genes. It consists of three core proteins: Lysine specific demethylase 1 (Lsd1/Kdm1a), Co-repressor of REST (CoREST) and Histone deacetylase 1/2 (HDAC1/2), as well as several other subunits (*10–17*). The CoREST complex is capable of repressing transcription through coordinated deacetylation and demethylation of histones (*18–22*). The Lsd1-CoREST interaction as well as the tertiary complex have been extensively examined, both structurally and biochemically, and CoREST has been shown to be required for demethylation of lysine 4 of histone H3 (K3K4) on nucleosome substrates by Lsd1 in mammals (*19, 21, 23–27*). The presence of this complex has been shown in mammals (*14*), *Drosophila melanogaster* (*28*) and *Caenorhabditis elegans* (*29, 30*) indicating it is at least conserved throughout Bilateria. The CoREST complex and CoREST proteins have been shown to play roles in the differentiation and homeostasis of various tissues, including the nervous system (*31–36*), epidermis (*37*), immune system (*38*) and the hematopoietic system (*39*) in mammals. In *Drosophila*, dLsd1 and CoRest play roles in spermatogenesis and follicle cell differentiation in the ovary (*28, 40, 41*) while in *C. elegans* homologs of both are involved in reprogramming of the zygotic genome after fertilization, something which is conserved in mice (*42–44*).

*Nematostella vectensis*, the starlet sea anemone, represents a particularly attractive system to investigate the evolution of animal development. It is a member of the phylum Cnidaria, the sister group to the bilaterian animals, and therefore possesses an informative phylogenetic position which, through comparative analysis, can unveil aspects of early animal evolution. In addition, a diverse array of experimental tools and resources is available for *Nematostella* (*45, 46*). Like most cnidarians, *Nematostella* has a simple adult body plan, known as a polyp, which resembles a tube with an opening at one end (oral) which serves as the mouth/anus and is surrounded by a ring of tentacles. Adults are dioecious and release sperm/eggs into the water column where fertilization occurs. Development proceeds through a hollow blastula which gastrulates via invagination to generate the two germ layers: ectoderm and endoderm. Following gastrulation, the animal develops into a free-swimming planula larva which eventually settles and metamorphoses into a primary polyp which will feed and grow to sexually maturity in approximately three months (*47–49*).

We have previously shown that the *Nematostella* ortholog of *Lsd1*, *NvLsd1*, in expressed ubiquitously throughout development but that its levels are developmentally regulated and are specifically high in differentiated neural cells relative to their progenitors. Using a mutant allele, we have also shown that loss of *NvLsd1* leads to a range of developmental abnormalities, the most pronounced of which is the almost total loss of differentiated cnidocytes, cnidarian specific neural cells (*50*). Here, by interrogating the interactome of NvLsd1 we found that the CoREST complex is conserved in *Nematostella*. We show that NvLsd1 and NvCoREST are expressed in precisely the same fashion throughout development. Using two mutant lines we show that loss of *NvCoREST* phenocopies loss of *NvLsd1* and that the CoREST complex is required for normal development and cnidocyte differentiation in *Nematostella*.

## Results

### The CoREST complex evolved early in animal evolution

Using a literature and homology-based approach we searched for core components of the CoREST complex in the genomes of a representative group of eukaryotes (see materials and methods and table S1). Lsd1 is an ancient protein present in plants, animals, and fungi (*51*). Similarly, class 1 HDACs, to which HDAC1/2 belongs, are found in all 3 aforementioned clades with a definitive branching giving rise to HDAC1/2 occurring before the emergence of animals as indicated by its presence in choanoflagellates, the closest extant relatives to animals (*52–54*). CoREST orthologs, on the other hand are only present in animals (Fig. 1A, table S1). Based on the presence of all three core components in non-bilaterian animals we hypothesized that the CoREST complex predates the cnidarian-bilaterian split. To test this, we used an unbiased approach to identify interactors of NvLsd1. We took advantage of our previously characterized line in which we have endogenously tagged NvLsd1 with eGFP using CRISPR-Cas9 (*50*). We used co-immunoprecipitation (Co-IP) with the GFP-Trap system coupled to liquid chromatography-mass spectrometry (LC-MS) to identify proteins which interact with NvLsd1 at planula stage. We found that both NvCoREST and NvHDAC1/2 were highly enriched in the NvLsd1-GFP sample (Fig 1, B and C and table S2). In addition, we found two other proteins NvHMG20a and NvPHF21a, putative *Nematostella* orthologs of HMG20A/B (iBRAF/BRAF35) and PHF21A (BHC80), which have also been identified as components of the vertebrate CoREST complex (*14, 55–57*) (Fig. 1, B and C). In the case of NvPHF21a we were unable to use a Student’s t-test as there was a missing value in one of the samples (i.e. making n=2 and therefore not suitable to perform a Student’s t-test with all valid values). We are, however, confident that this is a bona fide interactor because it has a very high fold change which is consistent among the two replicates for which we have values for both *NvLsd1^GFP^* and control Co-IPs (table S2) and as it was among the most highly enriched proteins in a pilot experiment (along with the other four proteins shown here) (Data S1) As CoREST is an essential, core component of the complex we decided to investigate NvCoREST further. We generated a custom antibody against amino acids 1-199 of NvCoREST (fig. S1A). This antibody recognizes two bands by western blot (Fig. S1B) which correspond approximately in size to two splice isoforms of *NvCoREST* which we can detect by PCR and which we have cloned and sequenced (Fig. S1C). IP and western blot showed that NvLsd1 interacts with both isoforms of NvCoREST (Fig. 1D). Together this data shows that the CoREST complex is indeed present in *Nematostella* and contains the same subunits as present in vertebrates.

**Fig. 1.**
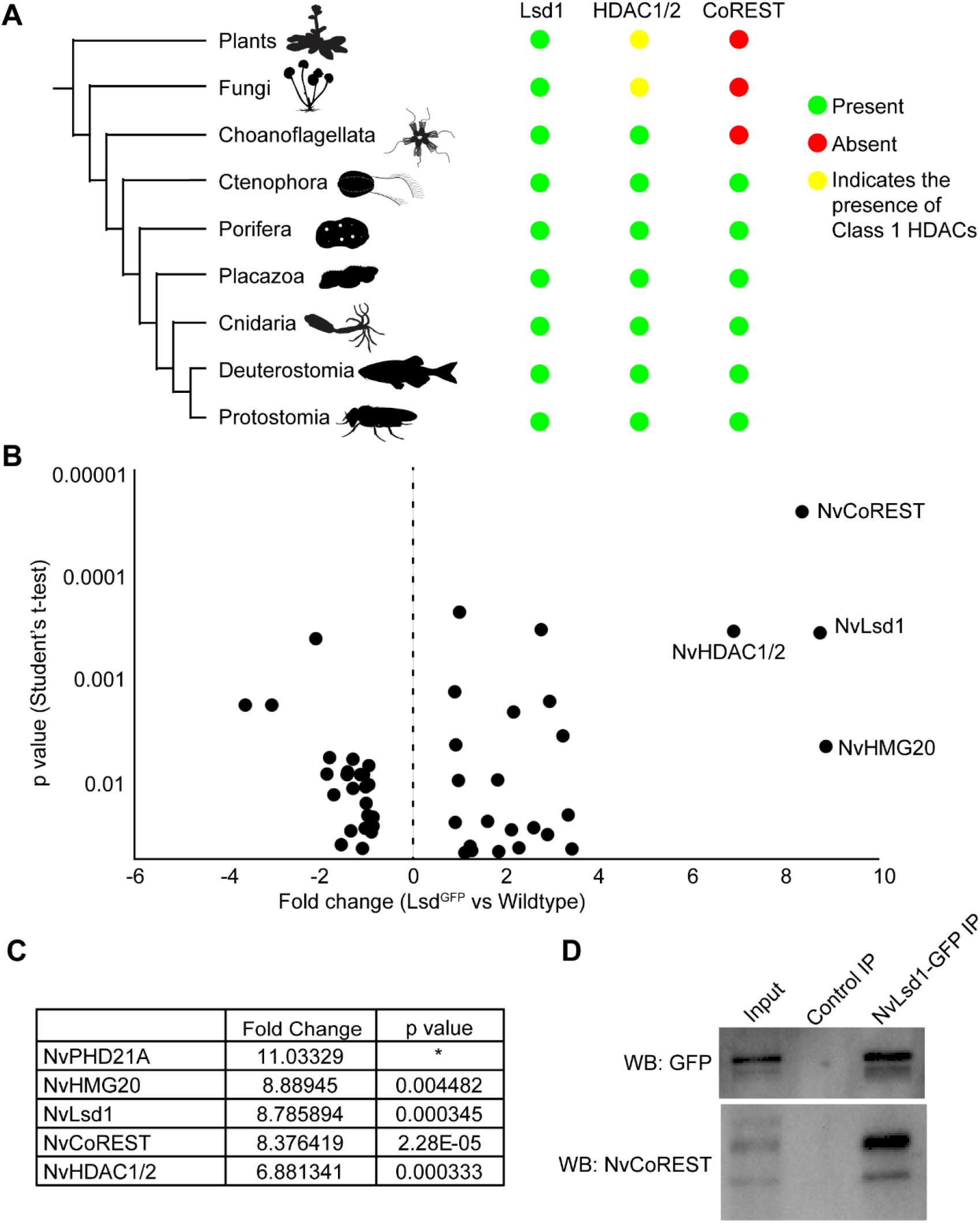
The CoREST complex is present in *Nematostella*. (**A**) Phylogenetic tree showing the presence/absence of Lsd1, HDAC1/2 and CoREST orthologs in the different groups. The tree on the left shows the relationships between the different clades analyzed. Green circles indicate the presence of orthologs within that group while red circles indicate their absence. A yellow circle is used to denote the presence of Class 1 HDACs as a definite HDAC1/2 orthologs are only seen in animals while the larger Class 1 group is more ancient. (**B**) Dot plot showing proteins derived from comparison of Co-IP with GFP-Trap beads from *NvLsd1^GFP^* and wild-type planula. Data is derived from three independent biological replicates. Only proteins with p-value <0.05 (Student’s t-test, two tailed) are shown. Fold change (FC) in the *NvLsd^GFP^* sample over wildtype is shown on the x-axis and p-value on the y-axis. The four proteins with the highest FC are annotated with names. (**C**) Table showing the fold change and p-value for the five most enriched proteins in the *NvLsd^GFP^* sample over wildtype. * We did not calculate a p-value for NvPHF21A due to a missing value in one of the control samples. (**D**) Co-IPs from *NvLsd1^GFP^* planula with either anti-GFP Trap beads or control agarose beads followed by western blot analysis with indicated antibodies. This was repeated 3 times independently with the same result.

### NvCoREST expression is high in differentiated neural cells

Having established that the CoREST complex is present in *Nematostella* we next wanted to understand how NvCoREST is expressed. Using immunofluorescence staining, we see that NvCoREST is ubiquitous and present in every nucleus at every stage studied except for mitotic nuclei from which is it excluded (Fig. 2, A to D and fig S2, A to C). The levels of NvCoREST are, however, heterogeneous and this heterogeneity appears gradually during development (fig. S2, A to C). Immunofluorescence staining in the NvLsd1-GFP line revealed that cells with higher levels of NvCoREST also have higher levels of NvLsd1-GFP (Fig. 1E). We have previously shown that NvLsd1 levels are high in differentiated neural cells but not their progenitors (*50*). We also show this here for NvCoREST using immunofluorescence staining in parallel with EdU labelling for proliferating cells/progenitors and by immunofluorescence staining in three different neural reporter lines: *NvNcol3*::mOrange2 which labels cnidocytes, a cnidarian specific neural cell type (*58*), *NvFoxQ2d*::mOrange which labels sensory cells (*59*) and *NvElav1*::mOrange which labels a large proportion of sensory cells and ganglion neurons (*60*). We find that NvCoREST is relatively low in proliferating cells and high in differentiated neural cells (Fig. 3, A to D). Together this shows that NvCoREST expression is fully overlapping with that of NvLsd1 and is high in differentiated cells of the nervous system relative to their progenitors.

**Fig. 2.**
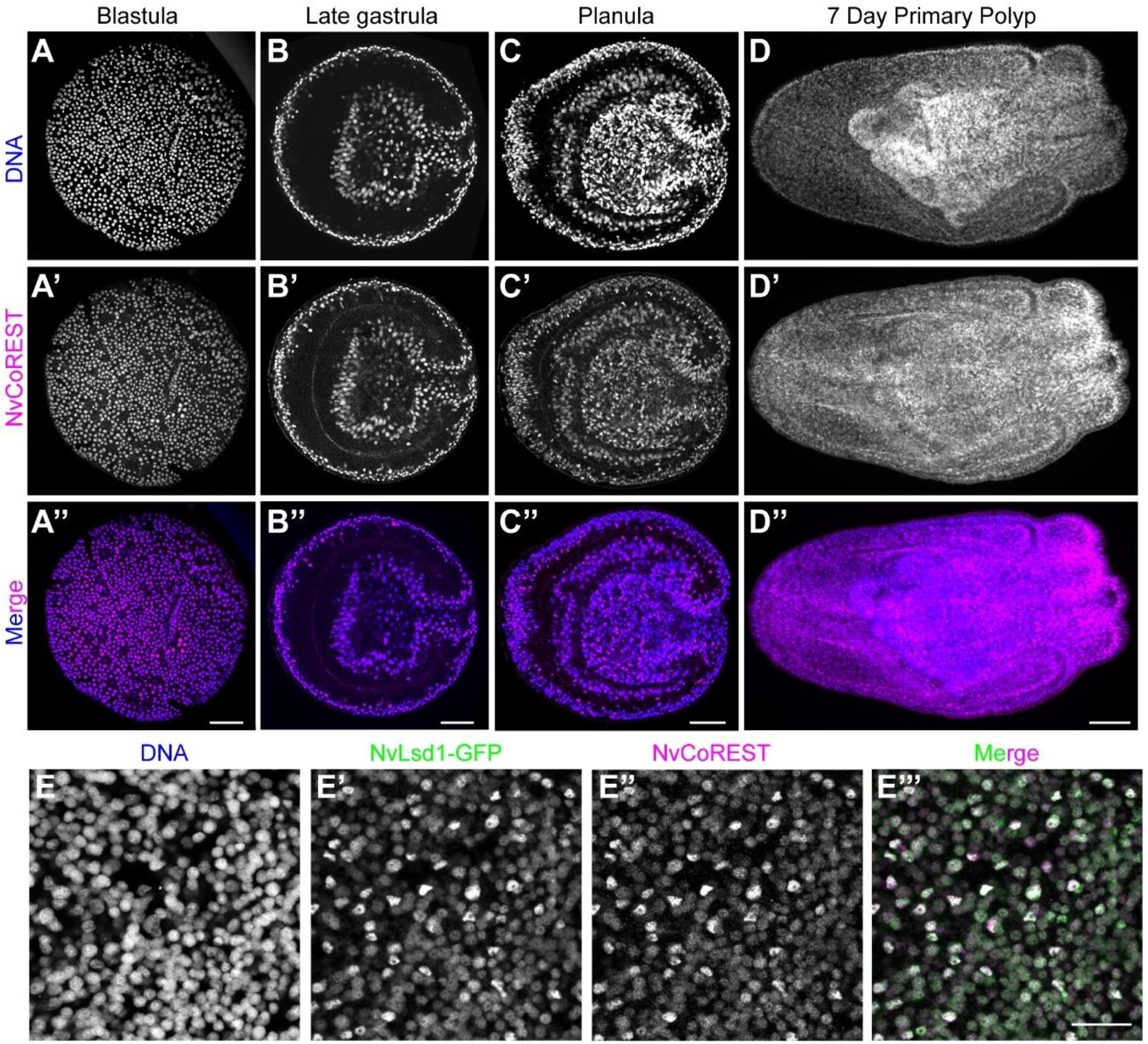
NvCoREST is ubiquitously expressed throughout development. (**A** to **D**) Confocal images of immunofluorescence staining showing NvCoREST localization throughout development. Stages are shown on top. (**E**) Close up of the ectoderm at planula stage showing colocalization of NvLsd1-GFP and NvCoREST. NvCoREST is shown in magenta, DNA in blue and NvLsd1-GFP in green. All staining’s were performed at least two times independently with a minimum of 10 embryos imaged per stage, per replicate with the same results. Scale bars: 50 μm (A to D). 20 μm (E).

**Fig. 3.**
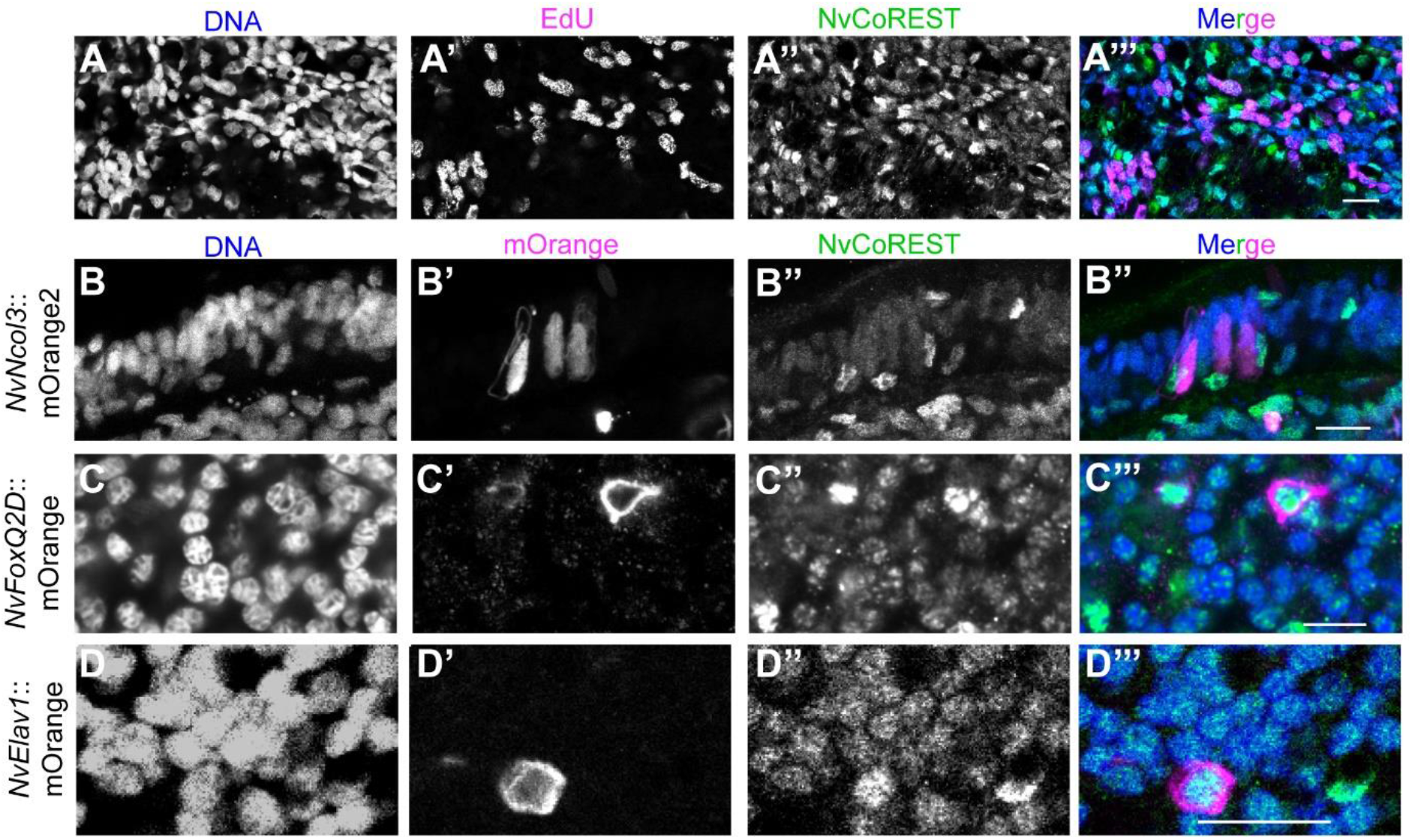
NvCoREST is low in proliferating cells and high in differentiated neural cells. (**A**) Confocal images of planula incubated with EdU for 30 minutes followed by immunofluorescence staining and Click-iT EdU detection. NvCoREST is shown in green, Click-iT EdU in magenta and DNA in Blue. (**B** to **D**) Confocal images of planula-stage transgenics stained for mOrange (magenta), NvCoREST (green) and DNA (blue) Transgenic lines used are indicated on the left. Stainings were performed two times independently with a minimum of 10 embryos imaged per genotype, per replicate with the same results. Scale bars: 10 μm

### *NvCoREST* is essential for *Nematostella* development

Next, to understand the function of *NvCoREST*, we generated two independent mutant lines using CRISPR-Cas9 targeting two different exons. Mutant 1 is an A-TGG substitution in exon 2 and Mutant 2 harbors the addition of a T in exon 3 (Fig. 4A). Both changes in the reading frame lead to premature stop codons and are predicted to generate early truncations of the NvCoREST protein (fig. S3A). In both mutant lines, we see that the homozygous mutants show a size defect at primary polyp stage (Fig. 4, B to E). We can separate the animals based on this phenotype and using sequencing we see that most animals sorted as having this phenotype are homozygous mutant (hereafter referred to as mutant) and, importantly, animals exhibiting a wild-type phenotype are never homozygous mutants (hereafter referred to as control) (Fig. 4, B to E). We also performed immunofluorescence staining for NvCoREST and we do not see any specific, nuclear staining in mutant animals from either line (fig. S3, B to E). In neither case were we able to find homozygous animals at juvenile or adult stage showing that they do not survive past this stage. A more detailed morphological analysis shows that despite the overall growth defect, mutant animals have metamorphosed and generated the normal structures expected to be present at this stage, i.e., four tentacles, pharynx, two primary mesenteries (Fig. 4, E to H). Overall, this data shows that *NvCoREST* is required for normal development in *Nematostella*.

**Fig. 4.**
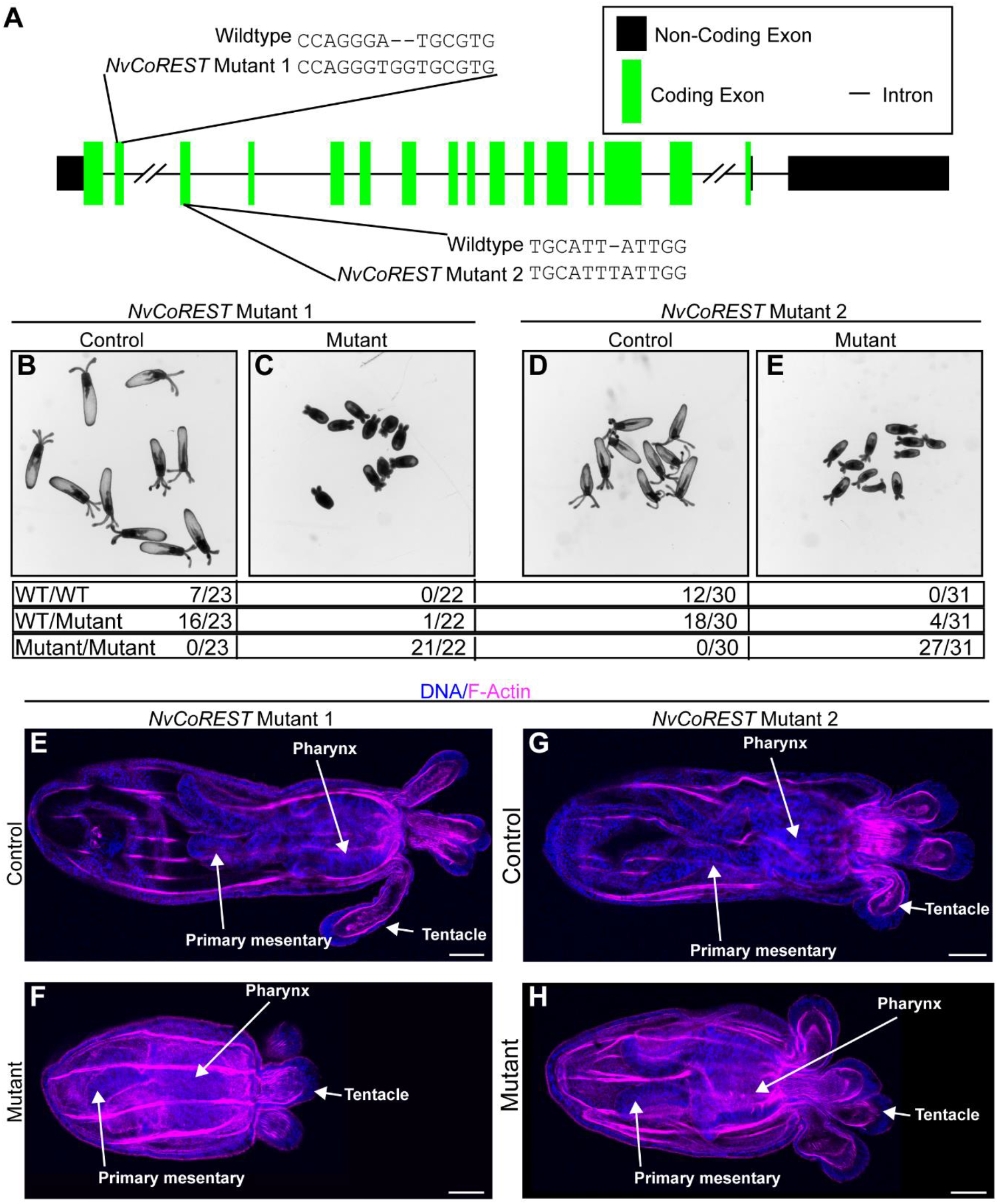
*NvCoREST* is required for normal development in *Nematostella*. (**A**) Schematic showing the intron-exon structure of *NvCoREST* and indicating both the position and sequence of mutations in *NvCoREST* Mutant 1 and Mutant 2 lines. (**B** to **E**) Live images of primary polyp stage animals generated from in-crosses of heterozygous *NvCoREST* Mutant 1 or Mutant 2 animals, separated based on size phenotype (indicated on top). Below the images is sequencing data showing the number of animals with the indicated genotype found via sequencing within these pools of animals. Numbers are combined data from 4 independent replicates. (**E** to **H**) Confocal images of representative animals from each phenotypic group stained with phalloidin for F-actin (magenta) and DNA (blue). Staining’s were performed there times, independently with a minimum of 10 embryos imaged per condition, per replicate with the same results. Scale bars: 50 μm

### Cnidocyte differentiation requires *NvCoREST*

We next looked at the effect of loss of *NvCoREST* on cnidocytes given that we have previously shown that *NvLsd1* is essential during cnidocyte differentiation. Cnidocytes are highly specialized, cnidarian-specific neural cells used for prey capture and defense and contain a specialized organelle, the cnidocyst, which contains a coiled thread that can be explosively discharged and acts like a harpoon (*61*). In both mutant lines we see that loss of *NvCoREST* leads to an almost complete loss of differentiated cnidocytes as determined using a protocol which utilizes DAPI to label the mature cnidocyst (Fig. 5, A to D) (*62, 63*). We also wanted to assess whether the reduction in cnidocyte number was due to a role for *NvCoREST* in specification or later differentiation of cnidocytes. To do so we performed immunofluorescence staining for NvNcol3, a protein found in the cnidocyst. The epitope recognized by this antibody, however, is only available prior to the maturation of the cnidocyst and thereby acts as a marker for earlier stages of cnidocyte differentiation (*64*). We found that there was abundant NvNcol3 staining in the *NvCoREST* mutants (fig. S4). The NvNcol3 staining, however, did not show the regular elongated capsules that are visible in the controls (arrows in fig. S4, A’ and A’’). This indicates that cnidocytes were still specified in the absence of *NvCoREST* but could not complete the differentiation process. In the case of *NvLsd1* we have previously shown that the requirement during cnidocyte differentiation is cell autonomous as the phenotype can be rescued by re-expression of *NvLsd1* using *NvPOU4* regulatory elements that drive expression predominantly in cnidocytes in the ectoderm (*50, 65*). We performed the same analysis here using an *NvPOU4::NvCoREST-mCherry* plasmid. We find that almost all cases (18/19) where we saw mosaic patches expressing NvCoREST-mCherry, we also see a rescue of the cnidocyte phenotype, something we do not see when we express NvHistone2B-mCherry as control (Fig. 5, E and F). Finally, we have also shown that loss of *NvLsd1* results in a phenotype in the *NvElav1*::mOrange^+^ nervous system characterized by a modest disorganization of the nerve net, the appearance of numerous mOrange^+^ puncta and expansion of the mOrange signal into surrounding epithelial cells. We have also looked here at the *NvElav1*::mOrange^+^ nervous system by crossing the *NvElav1::mOrange* transgene into the background of *NvCoREST* mutant 1. When we compare control and mutant animals, we do not see the same effect as in *NvLsd1* mutants (fig. S5). Together this shows that NvCoREST is essential for the post-mitotic differentiation of cnidocytes while being dispensable for the formation of the *NvElav1*::mOrange^+^ nervous system.

**Fig. 5.**
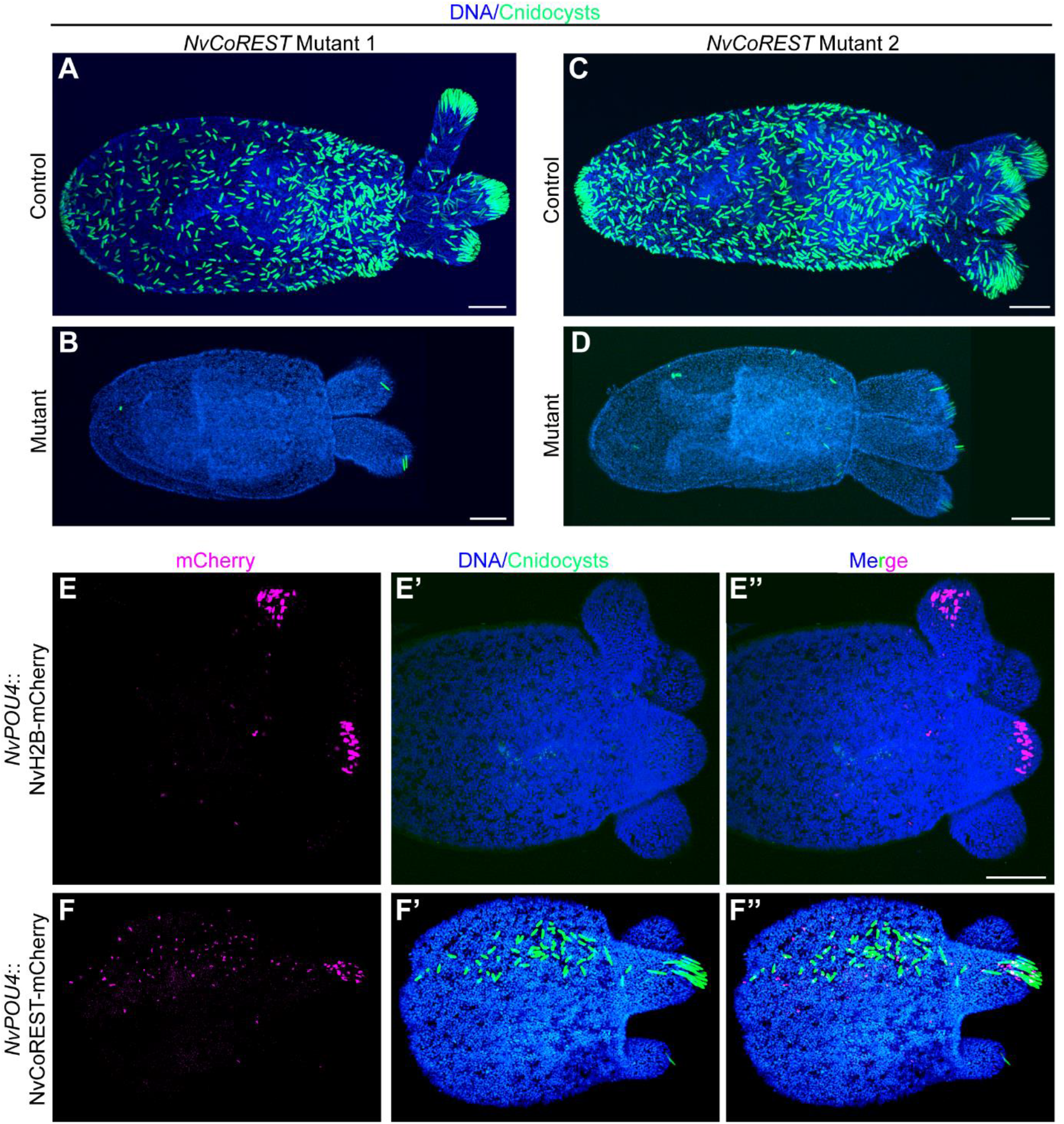
Loss of *NvCoREST* leads to a defect in cnidocyte differentiation. (**A** to **D**) Confocal images of control or mutant primary polyps from *NvCoREST* Mutant 1 or 2 showing cnidocysts (green) and DNA (Blue). Mutant line is shown on top and genotype to the left. Stainings were performed three times independently with the same results with a minimum of 10 embryos imaged per condition, per replicate. (**E** and **F**) Confocal images of immunofluorescence staining on mutant primary polyps from *NvCoREST* Mutant 1 line showing mosaic patches of NvPOU*4*::H2B-mCherry (E) or *NvPOU4*::NvCoREST-mCherry (F) expression stained with anti-DsRed antibody for mCherry (magenta), DAPI for cnidocysts (green) and DNA (Blue). Data was collected across two independent biological replicates and n=10 for *NvPOU4*::H2B-mCherry and n=19 for *NvPOU4*::NvCoREST-mCherry of which 10/10 and 18/19, respectively, had the observed phenotype. Scale bars: 50 μm.

## Discussion

We show here that CoREST is an animal-specific protein the evolution of which allowed the assembly of a novel chromatin-regulating complex early in animal evolution. We further show that the CoREST complex is present in *Nematostella* and that it is required during development and for the differentiation of cnidocytes. These observations provide a plausible explanation as to how the more ancient complex members, Lsd1 and HDAC1/2, may have been co-opted during evolution to play roles in development. The integration of these two chromatin modifiers into the CoREST complex thus may have facilitated the coordinated regulation of chromatin at specific genomic loci.

Our data point to a strong functional connection between NvLsd1 and NvCoREST. Not only do they interact but both proteins are also expressed in precisely the same manner and the phenotypes of loss of *NvCoREST* and *NvLsd1* are highly similar regarding the size of the animals as well as the loss of cnidocytes (*50*). We consider this as strong support for the hypothesis that most of the roles of NvLsd1 in *Nematostella* are mediated through the NvCoREST complex. A similar picture has also emerged in bilaterians where Lsd1 and CoREST interact, co-localize and indeed phenocopy each other in multiple systems (*14, 28–30, 40, 42*).

There are, however, differences between the two loss of function phenotypes. The size defect observed in *NvCoREST* mutants appears to be more severe that that seen in *NvLsd1* mutants, though it is possible that these differences result from the use of non-isogenic animals for generating the mutants. Secondly, we do not see the same effects in *NvCoREST* mutants on the *NvElav1*::mOrange^+^ nervous system as in *NvLsd1* mutants. This may indicate that some functions of *NvLsd1* are not mediated through the NvCoREST complex. This is interesting as roles outside of the CoREST complex have been assigned for NvLsd1 in mammals, particularly as a transcriptional activator e.g., in the mammalian nervous system (*66–69*). It will therefore be interesting in the future to more deeply analyze the function of *NvLsd1* in the *NvElav1*::mOrange^+^ nervous system and the development of tools for cell type-specific loss of function will greatly help in this regard.

Although CoREST is present in all animals it is possible that it first had a different function and only later became the scaffold of the CoREST complex as it exists in bilaterians. We have shown here that at least in the cnidarian-bilaterian ancestor this function was already present. Using our unbiased approach, we have shown that not only the three core components CoREST, Lsd1 and HDAC1/2 are present in the *Nematostella* CoREST complex, but the other two additional components of the vertebrate CoREST complex, PHF12A and HMG20, are also present. We searched for these genes in the *Drosophila* and *C. elegans* genomes using reciprocal blast searches but could not find them. This shows that the *Nematostella* complex is more similar to the vertebrate complex than those present in *Drosophila* and *C. elegans*. Furthermore, this suggests that the complex present in *Nematostella* and vertebrates is more similar to the ancestral complex while those present in *Drosophila* and *C. elegans* may be more derived. We have similarly recently shown that *Nematostella* contains a more vertebrate-like Polycomb Repressive Complex 1 (PRC1) repertoire at the level of presence/absence of complex specific components (*70*). Together this suggests anthozoan cnidarians and vertebrates may have retained more components of an ancestral machinery for chromatin regulation than other lineages and that *Nematostella* therefore is a useful model system in which to dissect fundamental aspects of the biology of such complexes.

## Materials and Methods

### Animal care and maintenance

*Nematostella* were maintained at 18-19°C in 1/3 filtered sea water [*Nematostella* medium (NM)], and spawned as described previously (*71*). Fertilized eggs were removed from their jelly packages by incubating in 3% cysteine in NM for 20 minutes followed by extensive washing in NM. Embryos were reared at 21°C and were fixed at 12 (blastula), 12 (blastula), 20 (early gastrula), 30 (late gastrula), 72 (planula), or at 13 days (primary polyp).

### Orthology search

The presence/absence of orthologs of Lsd1 and HDAC1/2 in different groups was extracted from existing literature (*51–54*). The presence or absence of CoREST homologs was determined using reciprocal Blast searches with the human CoREST1 sequency used as query. All genomes were searched using the NCBI database except *Mnemiopsis leidyi* for which we used the *Mnemiopsis* genome portal (https://research.nhgri.nih.gov/mnemiopsis/). A protein was considered an ortholog if it gave reciprocal blast hits with one of the human CoREST proteins and contained the domain architecture characteristic of CoREST proteins. The analyzed genomes are listed in supplementary table 1 and the accession numbers are given for genomes where CoREST orthologs were identified. In addition, choanoflagellates were not included in previous analyses and the presence of Lsd1 orthologs in this group was also assessed by reciprocal blast searches in the genomes of *Salpingoeca rosetta* and *Monosiga brevicollis.*

### Immunoprecipitation

Embryos from either wildtype or NvLsd1^GFP/GFP^ crosses were grown until planula stage. Approximately 50 μl of planula (volume of tissue without media) were used per IP. They were placed in lysis buffer (10 mM TricHCl pH 8, 150 mM NaCl, 2mM EDTA, 1% NP40, 10% glycerol) with cOmplete EDTA-free Protease Inhibitor Cocktail (Roche, 4693159001) and homogenized by passing through a 27G needle. Samples were then incubated on ice for 30 minutes and mixed approximately every five minutes by passing through the needle. Samples were then centrifuged at full speed for 15 minutes and 600ul of supernatant was used for IP. For each sample, 30 μl GFP-Trap Agarose or Binding Control Agarose Beads (Chromotek, gta-10 and bab-20) were washed once in dilution buffer (10mM tricHCl pH 7.5, 150 mM NaCl, 0.5mM EDTA) and then spun at 2500g for 2 minutes. The lysate was diluted with 900 μl dilution buffer and then added to the beads. This was incubated at 4°C for two hours rotating. Following this the beads were washed at least 6 times in 1 ml wash buffer (Dilution buffer + 0.5% NP40) for >10 minutes each at 4°C. In the final wash the beads were moved to a new tube. When protein was used for LC-MS analysis the wash buffer was removed and the beads were resuspended in 100 μl MilliQ H_2_O and frozen at −80°C until being processed further. In the case of western blotting the beads were incubated in 2X Laemmli sample buffer (0.1 M TrisHCl pH 6.8, 2% SDS, 20% Glycerol, 4% β-mercaptoethanol 0.02% Bromophenol blue) at 95°C, spun down and the supernatant was used for further analysis.

### Sample preparation for liquid chromatography-mass spectrometry (LC-MS)

Beads were thawed to room temperature (RT) and centrifuged at 2500g for 2 minutes and the H2O was removed. The beads were then resuspended in 40 μl trypsin buffer (50 mM Tris, 1mM CaCl_2_, pH8), 4 μl of 0.1 M DTT was added, and the samples were heated to 95°C for five minutes. The samples were then cooled to RT and 5 μl of 200 mM iodoacetamide was added, and the samples were incubated, shaking at RT for one hour. 0.8 μl of 0.1 M DTT was added to quench the remaining iodoacetamide, and samples were incubated shaking for 10 minutes. The pH was adjusted to approximately pH8 with 0.5 M Tris, 2 μg of Trypsin (Promega, V5111) was added to each sample and they were incubated shaking at 37°C overnight (o/n). Following this 5 μl of 10% trifluoroacetic acid was added to each sample and the peptide solutions were cleaned up with an Oasis HLB μElution plate (2 mg sorbent; Waters). Following elution samples were frozen at −80 and freeze dried.

### LC-MS analysis

Preliminary studies with samples containing 0.8 μg tryptic peptides dissolved in 2% acetonitrile (ACN) and 0.5% formic acid (FA) were injected into an Ultimate 3000 RSLC system coupled to a Q Exactive HF mass spectrometer (Thermo Scientific, Waltham, MA, USA). The MS1 resolution was 120 000 and the scan range 375-1500 m/z, AGC target was set to 3e^6^ and maximum injection time was 100 ms The intensity threshold was set at 5.0e^4^ and dynamic exclusion lasted for 20s. The MS/MS scans consisted of HCD with normalized collision energy at 28, quadrupole isolation window at 1.6 m/z and Orbitrap resolution at 15 000.

For the final experiments Control and *NvLsd1^GFP^* samples containing the same amount of peptide were analyzed in an Orbitrap Eclipse Tribrid mass spectrometer equipped with an EASY-IC/ETD/PTCR ion source and FAIMS Pro interface (Thermo Scientific, San Jose, CA, USA). The MS1 resolution and the scan range were set as above, AGC target was set to standard, maximum injection time was automatic and RF lens at 30%. The intensity threshold was also at 5.0e4 and dynamic exclusion lasted for 30s. The MS/MS scans consisted of HCD with collision energy at 30%, quadrupole isolation window at 4 m/z and Orbitrap resolution at 30 000. FAIMS was set up with the standard resolution mode and a total gas flow of 4.6 L/min. The CVs were set to −45 and −65 V.

### Statistical and bioinformatic analyses

The LC-Q Exactive raw files were searched in MaxQuant (version 1.6.14.0, Max Planck Institute for Biochemistry, Martinsread, Germany) (*72*) and the spectra were searched against the nveGenes.vienna database version 2008_02 (https://figshare.com/articles/dataset/Nematostella_vectensis_transcriptome_and_gene_models_v2_0/807696). The LC-Eclipse raw files were searched in Proteome Discoverer Software (version 2.5, Thermo Fisher Scientific, Bremen, Germany) using the SEQUEST HT database search engine with Percolator validation (FDR < 0.01), and against the uniprot-proteome UP000001593 database version 2021_02. Perseus (version 1.6.15.0, Max Planck Institute for Biochemistry) (*73*) was used to process and normalize the data. Proteins with three valid values in each group were selected for statistical comparisons using *t*-test. Proteins with *p*-values < 0.05 were considered to have significantly different abundance.

### Western blotting

Protein extraction was performed on Lsd^GFP^ or wild-type planula. Animals were placed in RIPA buffer (150 mM NaCl, 50 mM Tris pH8, 1% NP40, 0.5% DOC, 0.1% SDS) supplemented with cOmplete EDTA-free Protease Inhibitor Cocktail (Roche, 4693159001) and homogenized by passing through a 27G needle. Samples were incubated on ice for 30 minutes and mixed by passing through the needle every 5 minutes and centrifuged at full speed for 15 minutes at 4°C. The supernatant was kept and the protein concentration quantified using the Qubit™ Protein Assay (Invitrogen, Q33212). 30 μg of protein was used per lane, mixed 1:1 with 2X Laemmli sample buffer and boiled for 5 minutes before loading. For IP samples, beads were boiled in 2 volumes 2x Laemmli sample buffer for 5 minutes, spun down and the supernatant was loaded directly on to the gel. PageRuler™ Plus prestained protein ladder, 10 to 250 kDa (Thermo Scientific, 26619) was used. SDS PAGE was performed using 7.5% or 4-20% Mini-PROTEAN® TGX™ precast protein gels (BIO-RAD, 4561023/4561094) run in running buffer (25 mM Tris, 192 mM Glycine, 0.1% SDS) at 100 V for ~120 minutes. Transfer was performed using Trans-Blot Turbo Mini 0.2 μm PVDF Transfer Pack (BIO-RAD, 1704156) on a Trans-Blot Turbo transfer system (BIO-RAD) using the high molecular weight program. After transfer the membrane was washed in PBT (PBS + 0.1% Tween) several times and blocked with 5% milk powder in PBT (MPBT) at RT for 1 hour. The blots were incubated o/n at 4°C in 1° antibody in MPBT. The membranes were then washed several times in PBT and incubate in 2° antibody in MPBT at RT for 1 hour. Membranes were then washed several times in TBT and the signal was revealed using Clarify ECL substrate (BIO-RAD, 1705060) and imaged on a ChemiDoc XRS+ (BIO-RAD). The blots were then washed in PBT and blocked again in 5% MPBT for 1 hour at RT. They were then incubated o/n at 4°C with 1° antibody and processed as for the first antibody. Antibodies and dilutions are listed in table S4.

### Immunofluorescence

Animals at planula stage and older were anesthetized with MgCl2 and then killed quickly by adding a small volume (20-30 μl/ml) of 37% formaldehyde directly into the media. They were then fixed in ice cold 3.7% formaldehyde in PBTx [PBS(Phosphate Buffered Saline) + 0.2% Triton X-100] for 30-60 minutes (when staining for NvCoREST short fixations yield better staining) or for >60 minutes or o/n (for all other antibodies) at 4°C. Samples were washed >4 times in PBTx at RT, blocked in Block (3% BSA / 5% Goat serum in PBTx) for > 1 hour at RT and incubated in primary antibody diluted in Block o/n at 4°C. Samples were then washed extensively in PBTx (> 5 washes for 2 hours or more) at RT, blocked for 1 hour at RT in Block and incubated o/n or over the weekend in secondary antibody diluted in Block at 4°C. If Phalloidin staining was performed, Alexa Fluor™ 488 or 633 Phalloidin (Thermo Fisher Scientific, A12379/ A22284) was added here at 1:50-1:100. Samples were then incubated in Hoechst 33342 (Thermo Fisher Scientific, 62249) at 1:100 in PBTx for 1 hour at RT followed by extensive washing in PBTx (> 5 washes for 2 hours or more). Animals were mounted in ProLong™ Gold Antifade Mountant with DAPI (Thermo Fisher Scientific, P36935) and imaged on a Leica SP5 confocal microscope. Antibodies and dilutions are listed in table S4.

### EdU labelling

Animals used in Edu labelling experiments were incubated in 10 mM EdU in NM for the desired time and then treated for IF as described. After the final set of PBTx washes EdU incorporation was visualized using the Click-iT™ EdU Imaging Kit with Alexa Fluor™ 488 or 647 (Thermo Fisher Scientific, C10337/C10337) following the manufacturer’s protocol. Samples were mounted and imaged as for IF.

### DAPI staining for cnidocysts and counting

DAPI staining for cnidocysts was performed as previously published (*62, 63*) with slight modifications. Animals were processed as for IF with the addition of 10 mM EDTA to all solutions. Following the final PBTx wash, the samples were washed twice with MilliQ H_2_O and then incubated in 200 μg/ml DAPI in milliQ H_2_O o/n at RT. The samples were then washed once with MilliQ H_2_O, twice with PBTx with 10mM EDTA and mounted and imaged as for IF.

### CRISPR-Cas9 injections and genotyping

sgRNA was produced using a template generated by primer annealing. A PCR was set up containing 5 μl of each primer (100 mM) (One sgRNA specific and one generic), 2 μl dNTPs (10 mM each), 2 μl Q5 polymerase (NEB, M0491), 10 μl Q5 reaction buffer and 31 μl H2O with the following protocol: 98°C, 90 seconds; 55°C, 30 seconds; 72°C, 60 seconds. This was purified using a PCR clean up kit (Promega, A9281). The sgRNAs were synthesized using the MEGAscript™ T7 Transcription Kit (Invitrogen, AMB13345) including the DNase treatment and were precipitated by adding 1:1 LiCl (7.5 M) (Invitrogen AM9480) and incubating at −20°C for 30 minutes followed by centrifugation at full speed at 4°C for 15 minutes and extensive EtOH washes. The concentration was calculated using a Nanodrop. Primers are given in Table S3. *NvCoREST* mutants were produced similarly to previously published (*50, 74, 75*). Eggs were injected with a mix containing sgRNA (130 ng/μl), Cas9 (PNA Bio, CP01) (500 ng/μl) and 1:4 Dextran, Alexa Fluor™ 568 (Invitrogen, D22912) (200 ng/μl in 1.1 M KCl) that was incubated at 37°C for 5-10 minutes prior to injection. Injected animals were raised to sexual maturity and crossed to wildtypes. Individual F1 offspring were analyzed by PCR and sequencing to identify F0’s carrying the desired mutations. To extract genomic DNA, individual primary polyps were placed in tubes, the NM removed and 100% EtOH added. After 5 minutes this was removed, and the tubes were placed at 50°C for 45 minutes to allow the remaining EtOH to evaporate. 50 μl genomic extraction buffer (10 mM Tris pH8, 1 mM EDTA, 25 mM NaCl, 200 μg/μl ProteinaseK) was added to each and incubated at 50°C for 2 hours and 98°C for 15 minutes. 2 μl of this was used for PCR and sequencing. Primers are given in Table S3. Once an F0 carrier was identified, the remaining F1 offspring from that carrier were genotyped using a piece of tissue to generate a pool of F1 heterozygous animals which were then crossed and the offspring of these crosses were used in experiments.

### Cloning

For generating cDNA for PCR, RNA was extracted as previously published (*50*). The SuperScript™ III first-strand synthesis system (Invitrogen, 18080051) was used to generate cDNA. All PCRs were performed with Q5 polymerase and primers are listed Table S3. For cloning of the *NvCoREST* cDNA (Supplementary Fig. 2) the fragments were cloned using the CloneJET PCR Cloning Kit (Thermo Fisher Scientific, K1231). To generate the NvPOU4::NvCoREST-mCherry construct the backbone was amplified from the NvPOU4:: mCherry construct (*65*) and the CoREST open reading frame from cDNA, respectively. Assembly of the construct was done using the NEBuilder® HiFi DNA Assembly master mix (NEB, E2621)

### Transgenesis

In order to generate F0 mosaic transgenics we used I-SceI mediated transgenesis as previously described (*76*) with minor modifications. Eggs were injected with a mix containing: plasmid DNA (10 ng/μl), ISceI (1U/μl) (NEB, R0694), Dextran Alexa Fluor™ 568 (100 ng/μl), CutSmart buffer (1x). The mix was incubated for 30 minutes at 37°C before injection.

### Data and Material availability

All data necessary to reproduce the results are available in the main text or the supplementary materials. Novel reagents and animal lines are freely available upon request.

## Acknowledgements

We thank members of the Rentzsch lab for support and discussions throughout the project, Alexis Lanza for critical reading and comments on the manuscript and Eilen Myrvold and Lavina Jubek for excellent care of animals in the Sars Centre *Nematostella* facility. Research in FRs lab was funded by a grant from the University of Bergen and the Research Council of Norway (251185/F20) and by the Sars Centre core budget. Mass spectrometry-based proteomic analyses were performed by The Proteomics Unit (PROBE), Department of Biomedicine, University of Bergen. This facility is a member of the National Network of Advanced Proteomics Infrastructure (NAPI), which is funded by the Research Council of Norway INFRASTRUKTUR-program (project number: 295910).

## Author contributions

J.M.G designed and performed the experimental work, analyzed the data, conceptualized the study, generated the figures, and wrote the manuscript. M.H.V performed mass spectrometry and analyzed the data. F.R. conceptualized and supervised the study and wrote the manuscript. All authors edited the manuscript.

