## Supplementary material for "The chromatin-regulating CoREST complex is animal specific and essential for development in the cnidarian *Nematostella vectensis*": Supplemetary material

**This PDF file includes:**

Figs. S1 to S5

Tables S1 to S3

References

**Other Supplementary Materials for this manuscript include the following:**

Data S1

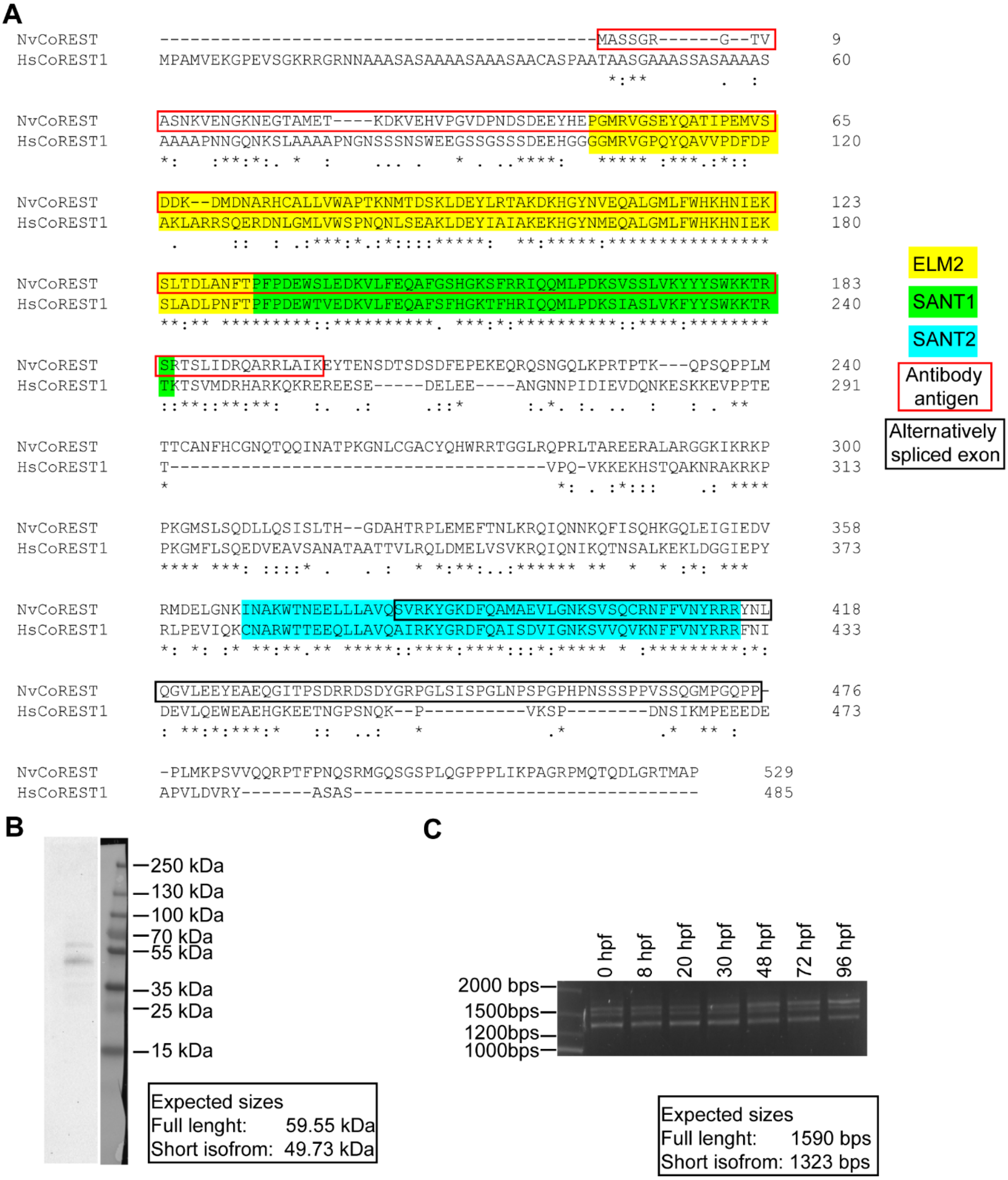


**Fig. S1. *NvCoREST* isoforms and antibody validation.** (**A**) Alignment of full length *Nematostella* CoREST with human CoREST1 (UniProt: Q9UKL0). Alignment was performed using Clustal Omega (*1*). Conserved domains are highlighted with coloured boxes; ELM2 in yellow, SANT1 in green and SANT2 in cyan. The portion of the protein used to generate the NvCoREST antibody is outlined with a red box. The alternatively spliced exon is outlined with a black box. (**B**) Western blot using anti-NvCoREST antibody showing two bands corresponding in size to the expected sizes of the full length and short isoform of NvCoREST, shown in the box at the bottom. Protein was extracted at planula stage. (**C**) PCR analysis using primers to amplify full length *NvCoREST* from cDNA from different developmental stages. The stage from which the cDNA was generated is shown on top measured in hours post fertilization (hpf). The expected sizes of full length and the short isoform of *NvCoREST* are shown in a box. Three bands are present; the highest and lowest correspond to the full length and short isoform of *NvCoREST*, respectively, and were successfully closed and sequenced. The middle band was never cloned and is presumably a PCR artifact, likely due to hybridization between the full length and short isoforms. Western blot and PCR analysis were carried out two times, independently with the same results.

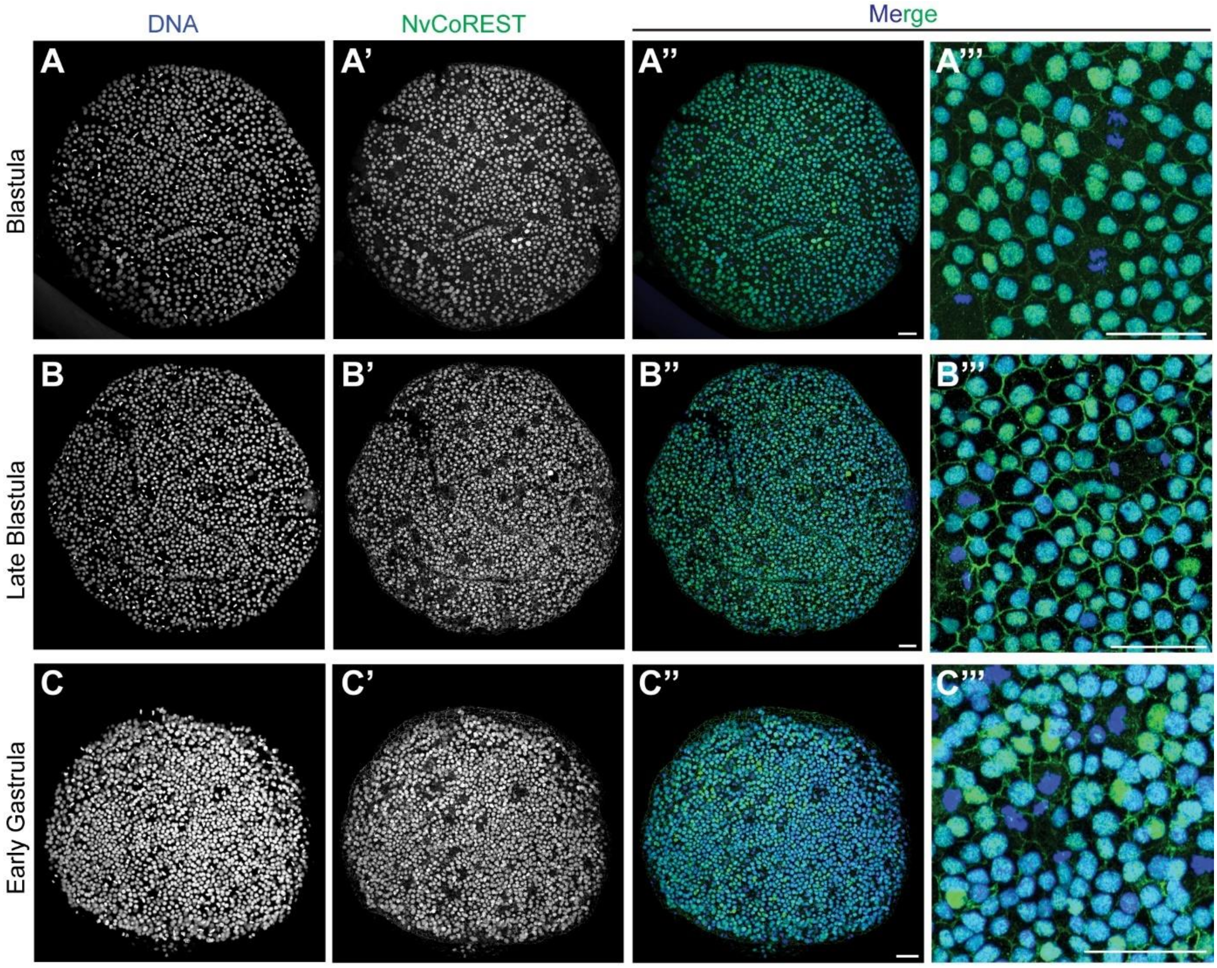


**Fig. S2. The heterogeneity in NvCoREST levels appears over developmental time.** (**A** to **C**) Confocal images of immunofluorescence staining performed on early embryos. Stages used are indicated to the left of the images. (A''' to C''') show close ups. NvCoREST is shown in green and DNA in blue. Staining's were performed two times independently with a minimum of 10 embryos imaged per replicate with the same results. Scale bars: 20 µm.

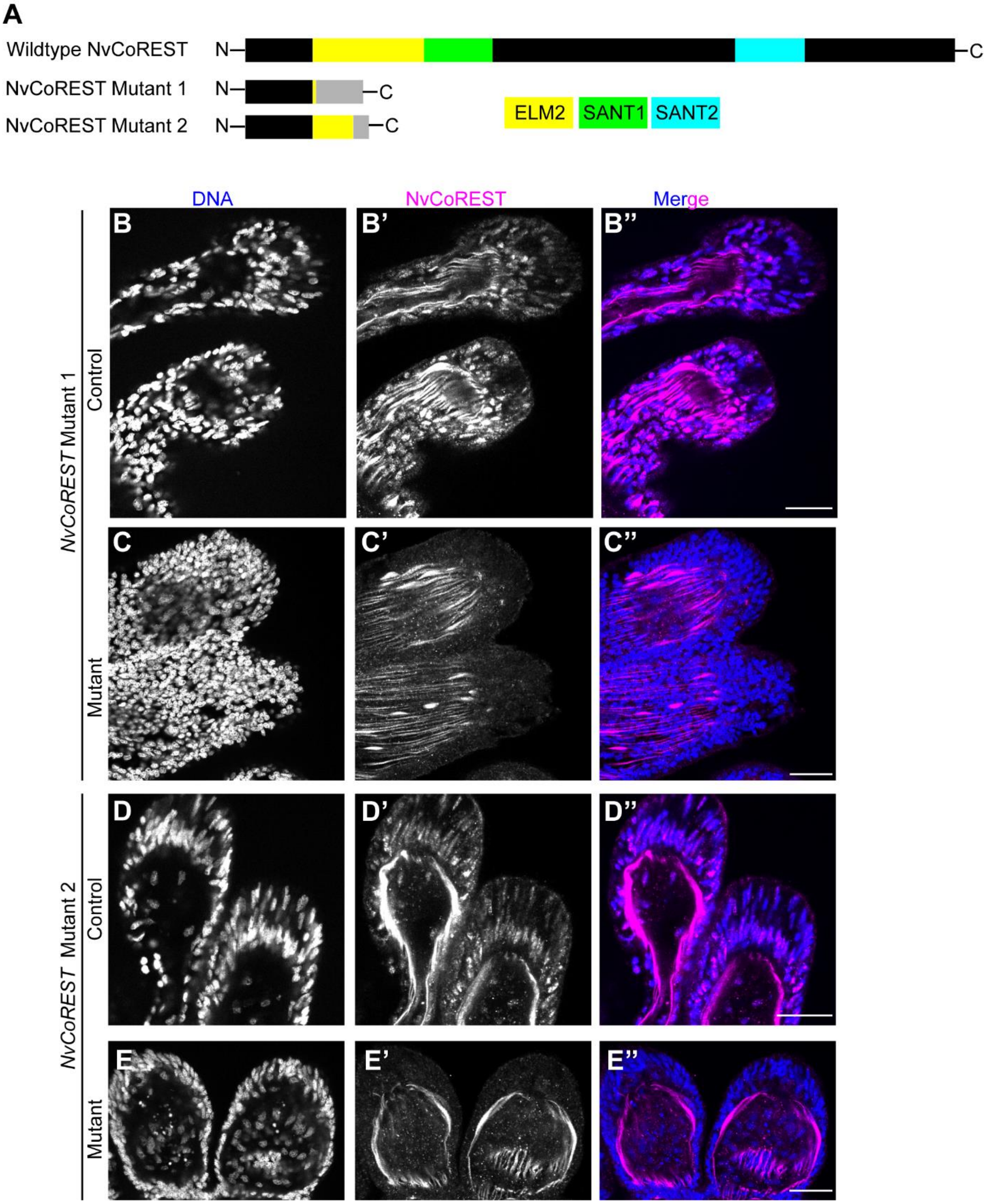


**Fig. S3. NvCoREST staining is absent in both *NvCoREST* mutant lines.** (**A**) Schematic representation of wildtype NvCoREST protein sequence alongside the predicted truncations in *NvCoREST* Mutant 1 and 2. Known domains are shown in colour, as indicated (**B** to **E**) Confocal images of immunofluorescence staining on mutant and control primary polyps from *NvCoREST* Mutant 1 or 2 lines stained for NvCoREST (Magenta) and DNA (Blue). Mutant line and genotype are shown to the left. Ubiquitous nuclear NvCoREST staining can be seen in control but is absent in mutant animals while non-specific staining of actin filaments can be seen in both. Stainings were performed two times independently with a minimum of 10 embryos imaged per genotype, per replicate with the same results. Scale bars: 20 µm.

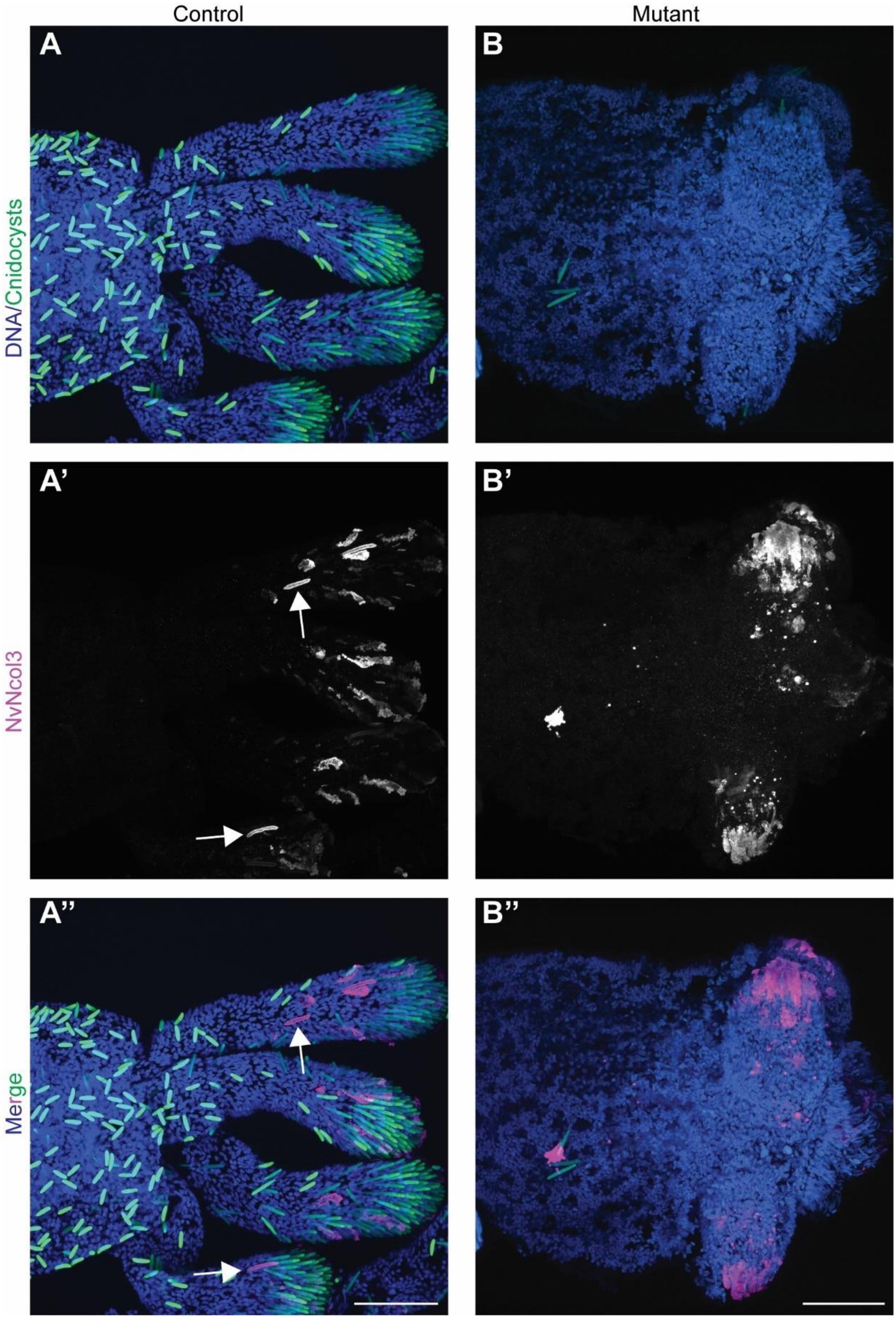


**Fig. S4. *NvCoREST* mutants still express NvNcol3.** (**A** and **B**) Confocal images of immunofluorescence staining on *NvCoREST* Mutant 1 primary polyps showing NvNcol3 in Magenta, DNA in blue and cnidocysts in Green. Arrows in A' and A'' indicate developing cnidocysts with normal morphology. The experiment was performed twice, independently and a minimum of 10 animals per genotype, per replicate were analyzed and showed the same result. Scale bars: 50 µm.

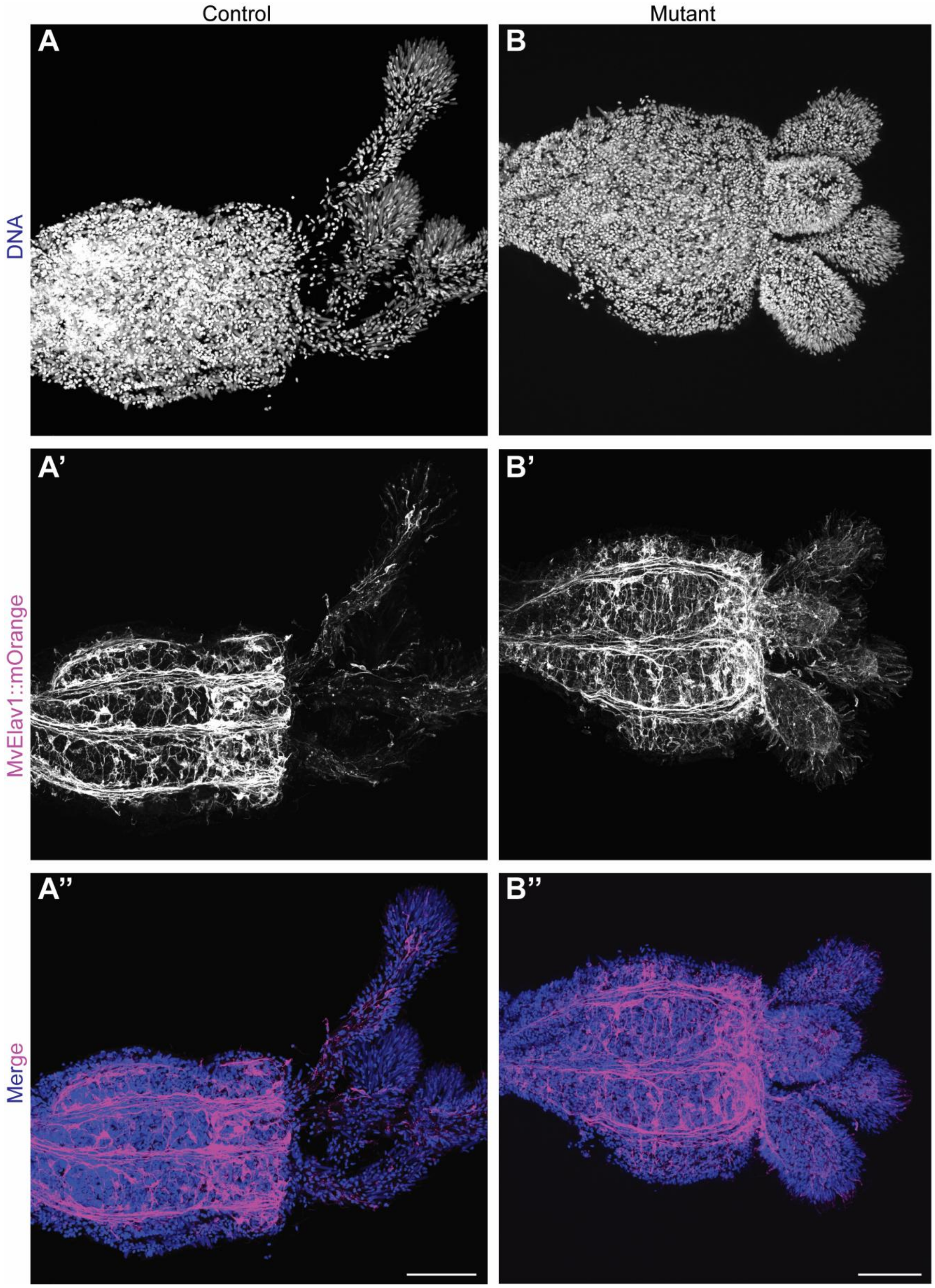


**Fig. S5. Loss of *NvCoREST* does not affect the *NvElav1*::mOrange$^+$ nervous system. (A** and **B**) Confocal images of immunofluorescence staining on control and mutant primary polyps showing DNA in blue and *NvElav1*::mOrange in magenta. Heterozygous *NvCoREST* mutant 1 animals were crossed to animals double heterozygous for *NvCoREST* mutant 1 and the *NvElav1*::*mOrange* transgene. The experiment was performed three times, independently and a minimum of 10 animals per genotype, per replicate were analyzed and showed the same result. Scale bars: 50 µm.

**Table S1 List of genomes used to search for CoREST othologs**

| Species name | CoREST (Presence/Absence) + Accession number |
|---|---|
| *Arabidopsis thaliana* | Absent |
| *Oryza sativa* | Absent |
| *Vitis vinifera* | Absent |
| *Selaginella moellendorffii* | Absent |
| *Spirogyra pratensis* | Absent |
| *Saccharomyces cerevisiae* | Absent |
| *Schizosaccharomyces pombe* | Absent |
| *Neurospora crassa* | Absent |
| *Capsaspora owczarzaki* | Absent |
| *Salpingoeca rosetta* | Absent |
| *Monosiga brevicollis* | Absent |
| *Amphimedon queenslandica* | Present (XM_019995969.1) |
| *Mnemiopsis leidyi* | Present (ML274412a) |
| *Nematostella vectensis* | Present (XM_032372883) |
| *Drosophila melanogaster* | Present (NP_001014754) |
| *Mus musculus* | Present (NP_001277209) |

### Table S2. Data from individual replicates of LC-MS experiments.

| Name | UniProt ID | NVE annotation | Control 1 | Control 2 | Control 3 | GFP 1 | GFP 2 | GFP 3 | Median Control | Median GFP | Fold Change |
|---|---|---|---|---|---|---|---|---|---|---|---|
| NVPHD21A | A7RH46 | NVE16417 | NaN | -4.68 | -4.00 | 6.19 | 5.61 | 6.66 | -4.34 | 6.19 | 11.03 |
| NvHMG20 | A7S5L8 | NVE23581 | -2.12 | -5.17 | -0.23 | 6.77 | 5.60 | 7.55 | -2.11 | 6.77 | 8.89 |
| NvLsd1 | A7S5A0 | NVE23413 | 2.05 | 2.74 | 0.09 | 10.94 | 10.84 | 10.46 | 2.05 | 10.84 | 8.78 |
| NvCoREST | A7SSF7 | NVE11839 | 0.63 | 0.30 | 0.71 | 9.00 | 8.36 | 9.51 | 0.63 | 9.01 | 8.38 |
| NvHDAC1/2 | A7RFA3 | NVE222 | 0.34 | 1.39 | 1.36 | 8.24 | 7.53 | 9.36 | 1.35 | 8.24 | 6.88 |

### Table S3. List of primers used in this study.

| Primer name | Sequence (5'-3') |
|---|---|
| NvCoREST-sgRNA1_Fwd (Used to generate NvCoREST Mutant 1) | TTCTAATACGACTCACTATAGGCGAGCCAACACGCATCCCGTTTTAGAGCTAGA |
| NvCoREST-sgRNA2_Fwd (Used to generate NvCoREST Mutant 1) | TTCTAATACGACTCACTATAGGCAAGACACTGTGCATTATGTTTTAGAGCTAGA |
| sgRNA_Rev (Genreic and used for both sgRNAs) | AAAAGCACCGACTCGGTGCCACTTTTTCAAGTTGATAACGGACTAGCCTTATTTTAACTTGCTATTTCTAGCTCTAAAAC |
| NvCoREST_Mutant1_Sequence_Fwd | GAATGAAGGAACTGCGATGGAGACTAAG |
| NvCoREST_Mutant1_Sequence_Rev | CCATCCGACACCATCTCGGG |
| NvCoREST_Mutant2_Sequence_Fwd | GACGCTTTCCACTTTATGCTTTCCC |
| NvCoREST_Mutant2_Sequence_Rev | GCTTACATTTACTATCTGTCATATTCTTGGTAGG |
| NvCoREST_Full-Lenght_Fwd | ATGGCTTCTAGCGGCCG |
| NvCoREST_Full-Lenght_Rev (Used for cloning the gene and RT-PCR) | TCAAGGTGCCATGGTCC |
| NvCoREST_NoSTOP_Rev (Used to amplify NvCoREST to clone in frame with mCherry | AGGTGCCATGGTCCTACCAAG |
| NvPOU4_Vector_Fwd | CACAGGACCTTGGTAGGACCATGGCACCTGCGGGCGGCGGCGGCAGC |
| NvPOU4_Vector_Rev | GCCACGGTTCCTCGGCCGCTAGAAGCCATCGTGGAGCACTCAGCACCAAC |

### Table S4 List of antibodies and dilutions

| Name | Company | Catalogue number | Concentration (IF) | Concentration (Western) |
|---|---|---|---|---|
| Rabbit anti-DsRed | Clontech | 632496 | 1:100 | |
| Mouse anti-mCherry | Clontech | 632543 | 1:100 | |
| Chicken anti-GFP | Kerafast | EMU101 | 1:100 | |
| Rabbit anti-GFP | Abcam | Ab290 | | 1:20,000 |
| Rabbit anti-NvCoREST | Custom | n/a | 1:100 | 1:10,000 |
| Goat anti-rabbit Alexa 488 | Life Technologies | A11008 | 1:250 | |
| Goat anti-rabbit Alexa 568 | Life Technologies | A11011 | 1:250 | |
| Goat anti-mouse Alexa 488 | Life Technologies | A11001 | 1:250 | |
| Goat anti-mouse Alexa 568 | Life Technologies | A11004 | 1:250 | |
| Goat anti-chicken Alexa 633 | Life Technologies | A21103 | 1:250 | |
| Goat anti-chicken Alexa 488 | Life Technologies | A11039 | 1:250 | |
| Goat Anti-Rabbit (HRP) | Abcam | Ab97051 | | 1:10,000 |

### Data S1. (Separate file). Data from LC-MS experiments.
